# A Mathematical Model of the Effect of Natural Selection on Adaptation Forms that Implemented by disruptive coloration of *Taurotragus oryx*

**DOI:** 10.1101/368084

**Authors:** Yu. G. Bespalov, K. V. Nosov, P. S. Kabalyants

## Abstract

In the study, results of mathematical modelling of the influence of natural selection on performance of various forms of animal adaptation to habitat conditions are presented. For a formalized description of the subject of study, we used a new class of mathematical models— discrete models of dynamical systems. Sets of strategies of protective coloration of antelopes Taurotragus oryx are the subject of a formalized description. Various combinations of brightness of green and red components of gray-brown non-uniform protective coloration of different parts of the silhouette of these animals were considered as such strategies. The sets based on the material of digital pictures of the two groups of Taurotragus oryx were compared. The first group includes antelopes Taurotragus oryx from Serengeti National Park (Tanzania) exposed to natural selection. The second group includes Taurotragus oryx, actually domesticated in Askania-Nova reserve (Ukraine), for which natural selection is not active. The sets of above mention strategies-combinations, modelled for the two groups, were compared by the numbers of unique combinations of values of brightness of red and green colours, as well as combinations with closest values of these brightness. The adaptive role of combinations with different values of red and green colours was identified with the role of idioadaptations. The adaptive role of combinations with equal values of red and green colours was identified with a more wide performance of aromorphoses. In this connection, the notions “quasi-idioadaptation” and “quasi-aromorphosis” were introduced in the paper.

It is assumed that both quasi-aromorphoses and quasi-idiadaptations, in certain conditions, contribute to the destruction of an integral visual perception of the silhouette of an animal against a many-coloured background of vegetation. At that, assumed that an adaptation function of a quasi-aromorphosis can be implemented in a wider range of colorimetric parameters of a plant background. The results of modelling indicate that the coloration of Taurotragus oryx from Serengeti is characterized by a larger set of quasi-adaptations than coloration of Taurotragus oryx from Askania-Nova. In the coloration of the latter, there is no quasi-aromorphosis with maximum values of brightness of both red and green components. But there exists a quasi-aromorphosis in the coloration of Taurotragus oryx from Serengeti. Such results of mathematical modelling correspond to prevailing ideas about the influence of natural selection on the character of adaptive reactions of living beings.

## Introduction

One of the fundamental concepts of the evolutionary theory is an influence of natural selection on the character of adaptation forms (AFs) of living organisms. In this case, one can distinguish AFs, which have an adaptive meaning in a relatively narrow and in a fairly wide range of values of habitat parameters. For these AFs, the terms “idioadaptation” and “aromorphosis” were suggested [1].

The concepts of “idioadaptation” and “aromorphosis” are not currently commonly used. To denote the AFs with different degrees of universality, they use different terms. So in connection with this, in the framework of this work, similar to with idioadaptation and aromorphosis, the working terms “quasi-idioadaptation”(QIA, Quasi-IdioAdaptation) and “quasi-arorphosis” (QAM, Quasi-AroMorphosis) will be used.

In particular, different combinations of the values of certain parameters that provide an adaptation of a living organism to certain combinations of values of environmental parameters may correspond to different AFs. In the framework of the paper we are talking about a ratio of brightness of the red and green components of the protective disruptive animal coloration (*Taurotragus oryx*). Different ratios of mentioned parameters make it possible to realize the protective effect of coloration, which disrupts the visible contour of bodies of these antelopes against a background of vegetation, on different parts of which different ratios of green chlorophyll, red, orange and yellow phytopigments are present at different times during a growing season. In this case, the maximum diversity of these combinations (for example, the entire range of shades on different sites of vegetation at different time periods with predominance of red or green) will contribute to the protective effect of the QIA type. A combination, which is as close as possible to any of the indicated shades (in a rather simple case it can be a brown colour formed by mixing in equal proportions of red and green), will correspond to a protective effect of the QAM type.

The presence of these protective effects of QIA and / or QAM type allows us to study certain aspects of the problem of the influence of biodiversity on the effectiveness of forms of adaptation of biological systems. A stability can be considered as a criterion of efficiency (in our case study, the efficiency of antelopes’ protecting from predators in one way or another affects the stability of their population).

In the case of the implementation of an AF of the QIA type, the number of combinations of values of the system’s components plays the role of the determining aspect of an aspect of biodiversity. In the case of the QAM type, the evenness of the values of these components can be considered as an aspect of biodiversity.

In [2], it was proposed to use the number of possible strategies for performance of the system as an indicator of biodiversity. It is assumed that different combinations of values of the system’s components correspond to these strategies. These combinations, in particular, can be observed in an idealized trajectory of the system (ITS), reflecting dynamism of changes in system’s states determined by the structure of relationships between the components. Such a cycle can be observed, in particular, during the growing season for colorimetric parameters (CPs) of vegetation related to different content of green chlorophyll, and red, orange and yellow phytopigments. A certain set of animal coloration hues with different ratios of brightness of red and green may corresponds to the ITS that reflects this cycle. This set provides a protective effect of the dismemberment of a body contour against a vegetal background at various points of time and space. (The ratio of red and green in the colour of a plant background at each such point must match the same colour ratio of animal coloration at least for one spot, the size and location of which on a body is capable to break its silhouette). This set of colours can be represented by a matrix similar to an ITS’s matrix, but, unlike ITS, this matrix does not reflect any dynamics. For this matrix, the working term “idealized pseudo-trajectory of the system” (IPTS) was suggested. A set of columns of an ITS and IPTS with different combinations of component values may represent a diversity of above said strategies of system performance (e.g., the performance of protective coloration of an animal, which body has spots with different combinations of shades of red and green). With the help of a new class of mathematical models developed with authors’ participation — discrete models of dynamic systems (DMDS), which can be applied for a formalized description of performance of systems of different nature [2, 3, 4, 5], ITSs and IPTSs can be built on the basis of data of relatively small size with lacunae.

This work aimed at modeling, with evolving the DMDS models, the influence of natural selection on the nature of the protective disruptive coloration of an animal (*Taurotragus oryx*) with use of adaptation forms of the QIA and QAM types.

## Materials and methods

To study the effect of natural selection on the nature of the protective effect of animal disruptive coloration, we performed a comparative analysis of IPTS’s fragments, taken from [6], built with use of DMDS. A modelling in this work were performed with use of Spearman correlations and the approach based on the Liebig law of minimum [3]. The data of the IPTS reflect a distribution of different combinations of the values (in conditional scores) of colorimetric parameters of the two groups of *Taurotragus oryx*. The images used for study are freely available online from numerous resources, for example, at https://wikivisually.com. The results were obtained with use of the freely available Matlab package (https://sites.google.com/site/improctoolkit/) based on analysis of the RGB model of digital images. We are talking on the following two groups of these antelopes: practically domesticated in the Askania-Nova Reserve (Ukraine) and those that live essentially in wild conditions under the influence of natural selection factors in the Serengeti National Park (Tanzania)

## Results and Discussion

A comparative analysis of IPTSs for coloration of Serengeti’s antelopes that live under conditions of natural selection factors (including suppression by predators), on one hand, and practically domesticated Askania-Nova’s antelopes, where natural selection factors are not active (at least there is no predators’ suppression, for protection from which disruptive coloration serves).

The two IPTSs have the same (8) number of different combinations of the following four colorimetric parameters:

R/(R+G+B): corresponds to the red coloration component of antelopes, allowing them to camouflage against vegetation plots with a predominance of red, orange and yellow phytopigments;

G/(R+G+B): corresponds to the green coloration component of antelopes, allowing them to camouflage against vegetation plots with predominated chlorophyll;

G/(R+G): high values of this CP correspond to vegetation areas with high productivity;

R/G: high values of this CP correspond to vegetation areas with stabilized bioproduction processes.

For the aims of this study, a comparative analysis of IPTS rows is important, as it gives an idea about the distribution of combinations of red and green components of antelope coloration (colorimetric parameters R/(R+G+B) and G/(R+G+B)). This distribution is presented, for Serengeti’s antelopes, in Table 1, for Askania-Nova antelopes — in Table 2.

**Table 1.**
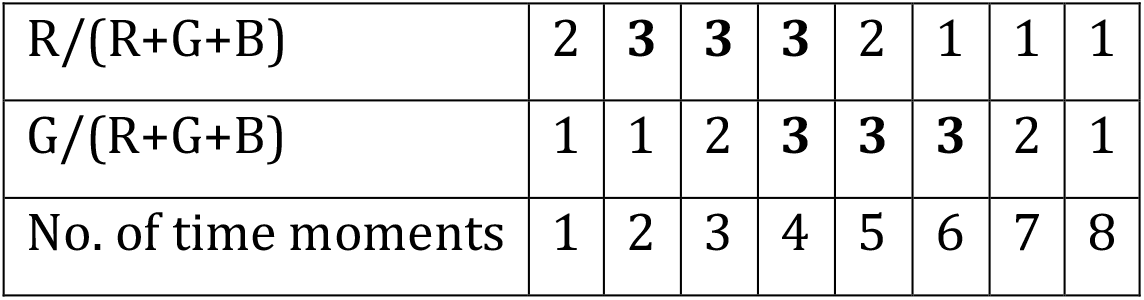
Fragment of the IPTS, built on the material of antelope coloration of Serengetti. Rows: the values of colorimetric parameters in conditional scores (1 - low, 2 - medium, 3 - high). Columns: conditional time steps. The maximum values of the colorimetric parameters for a given IPTS are in bold.

|  |  |  |  |  |  |  |  |  |
| --- | --- | --- | --- | --- | --- | --- | --- | --- |
| $R/(R+G+B)$ | 2 | **3** | **3** | **3** | 2 | 1 | 1 | 1 |
| $G/(R+G+B)$ | 1 | 1 | 2 | **3** | **3** | **3** | 2 | 1 |
| No. of time moments | 1 | 2 | 3 | 4 | 5 | 6 | 7 | 8 |

**Table 2.** Fragment of the IPTS, built on the material of antelopes coloration in Askania-Nova. Notation similar to Table 4.

|  |  |  |  |  |  |  |  |  |
| --- | --- | --- | --- | --- | --- | --- | --- | --- |
| $R/(R+G+B)$ | 1 | 1 | 1 | 2 | 2 | 2 | 2 | 1 |
| $G/(R+G+B)$ | 1 | 2 | 3 | 3 | 3 | 2 | 1 | 1 |
| No. of time moments | 1 | 2 | 3 | 4 | 5 | 6 | 7 | 8 |

We see that Table 1 has more unique combinations of values of the colorimetric parameters reflecting the ratio of red to green components of antelope coloration than Table 2 (8 vs. 6).

This can be interpreted as the influence of a weakening (or even disappearance) of the suppression of natural selection on the diversity of AFs of the QIA type.

Additionally, Table 1 contains a combination with equal maximum (three scores) values of the parameters R/(R+G+B) and G/(R+G+B), but Table 2 does not contain such the combination. The combination featured by equal (high, medium or low) values of brightness and, correspondingly, high values of evenness of brightness of red and green, has a certain range of conditions for the implementation of masking effect, since it can enable the fusion of a spot on the animal’s body in vegetation areas accompanied with deviation of a ratio of colorimetric parameters in both green and red side. Therefore, the form of adaptation implemented by such a combination can be classified as QAM. For Serengeti’s antelopes, this form of adaptation is manifested for high (at 4th time step) and low (8th time step) values of the colorimetric parameters R/(R+G+B) and G/(R+G+B). The combination of high values of these parameters with a high rate of their evenness for *Taurotragus oryx* in wild has a significant adaptive effect because it provides masking from predators in areas with high values of biomass of grass eaten by antelopes.

For actually domesticated antelopes in Askaniya-Nova provided with additional food and protection from predators, this adaptive effect neither matters nor does it affect the nature of protective coloration. Hence, the presence in Table 2 of combinations of values of the colorimetric parameters R/(R+G+B) and G/(R+G+B) corresponding to AFs of the QAM type that have medium (6th time step) or low (1st and 8th steps) scores is quite understandable.

The results of mathematical modeling presented in Table 1 and Table 2 correspond to generally recognized ideas about formation of certain forms of adaptation under the influence of natural selection. This effect leads to an increase in the number of combinations of values of different system’s components (in our case, the brightness of green and red colors in animal coloration). In work [2], these combinations were interpreted as strategies of performance of the system. In this paper, we interpret them as strategies of implementation of AFs of the QIA type. The influence of natural selection leads also to appearance of adaptation forms of the QAM type having high values of system’s components and high evenness of these values (in the considered case, parameters R/(R+G+B) and G/(R+G+B)).

Optimal structures, for certain systems in some conditions, of relations between AFs of the QIA and QAM type are to be the main subject of further research. It makes sense to study dependences of such structures on the diversity of conditions to which the systems should adapt (optimal number of QIAs), a permissible degree of compliance or non-compliance of QAM with a specific set of these conditions.

In this context, the role of such aspects of biodiversity as the number of strategies in system performance and the degree of evenness in combinations of values of system’s components that correspond to these strategies can be studied. At that, one can also interpret these combinations-strategies as different ways of using system resources, as suggested in [6].

We can conclude, that results obtained in this paper create certain opportunities for conducting specified studies.

